# CNC Knitting Micro-Resolution Mosquito Bite Blocking Textiles

**DOI:** 10.1101/2023.04.21.537869

**Authors:** Bryan Holt, Kyle Oswalt, Alexa England, Richard Murphy, Isabella Owens, Micaela Finney, Natalie Wong, Sushil Adhikari, James McCann, John Beckmann

## Abstract

Mosquitoes and other biting arthropods transmit diseases worldwide, causing over 700,000 deaths each year, and costing about 3 billion annually for *Aedes* species alone. These insects also pose a significant threat to agricultural animals. While clothing could provide a simple solution to vector-borne diseases, modern textiles do not effectively block mosquito bites. To address this issue, we have designed three micro-resolution knitted structures, with five adjustable parameters, that can block bites. These designs were integrated into a computer numerical control knitting robot for mass production of bite-blocking garments with minimal human labor. We then quantified the comfort of blocking garments. Our knits enable individuals to protect themselves from insects amidst their day-to-day activities without impacting the environment.

**One Sentence Summary:** We create micro-resolution mosquito bite blocking knits produced by robotic manufacturing to protect humans against vector-borne disease.

## Main Text

Mosquito bites transmit diverse pathogens including viruses, unicellular organisms, and even multicellular nematodes ^1^. More than half a million people die of malaria each year; most are young children ^2^. Controlling mosquito populations and these diseases remain a global problem ^3^. Mosquito-borne diseases can be epidemic and spread rapidly with changes in agriculture and migration ^4^. Humans who labor outside in tropical climates are at highest risk. Mosquito populations, in part, are controlled by insecticides which promote resistance and are detrimental to the environment ^5,6^. Recent advances in mosquito genetics and biological control can lower mosquito populations without insecticides, but still fundamentally alter earth’s ecology in safe, modest ways ^7–14^. Textiles have always been a pragmatic deterrent of mosquito-borne disease in the form of bed nets ^15^. Furthermore, recent research reported constructions of mosquito bite blocking textiles ^16^. Although variables controlling blocking in combination with comfort have not been reported. Importantly, these textile applications without impregnated insecticides have zero negative side effects. Bizarrely, we found that modern clothing doesn’t stop mosquito proboscises; some clothing is worse than being stark naked if it prevents perception of feeling a mosquito land. Popular form-fitting athletic “heat-gear” exacerbates the problem and does not block bites.

Female mosquitos feed with piercing/sucking proboscises (**Fig 1A**). The proboscis has an outer labium (Lab). At feeding, the labium retracts exposing the fascicle, which is a repertoire of six serrated blades and microneedles bound together by liquid surface tension (**Fig 1B**) ^17–19^. The labrum (Lm) is a beveled needle which pierces and draws blood. Adjacent to the labrum are paired mandibular (Md) and maxillary (Mx) stylets. Maxillary stylets saw skin at a vibrational frequency of 30 Hz to reduce the force needed to puncture ^20^. The flexible fascicle can bend at 90° angles, is innervated and controlled by delicate musculature ^18,21^. The measurements of the proboscis in *Aedes aegpyti* (the yellow fever mosquito) are 2.32 mm long and 60 μm wide. The labrum is 25μm in diameter ^17,18,22–24^. Needless to say, designing clothing to mechanically block the mosquito fascicle is tough engineering.

**Figure 1.**
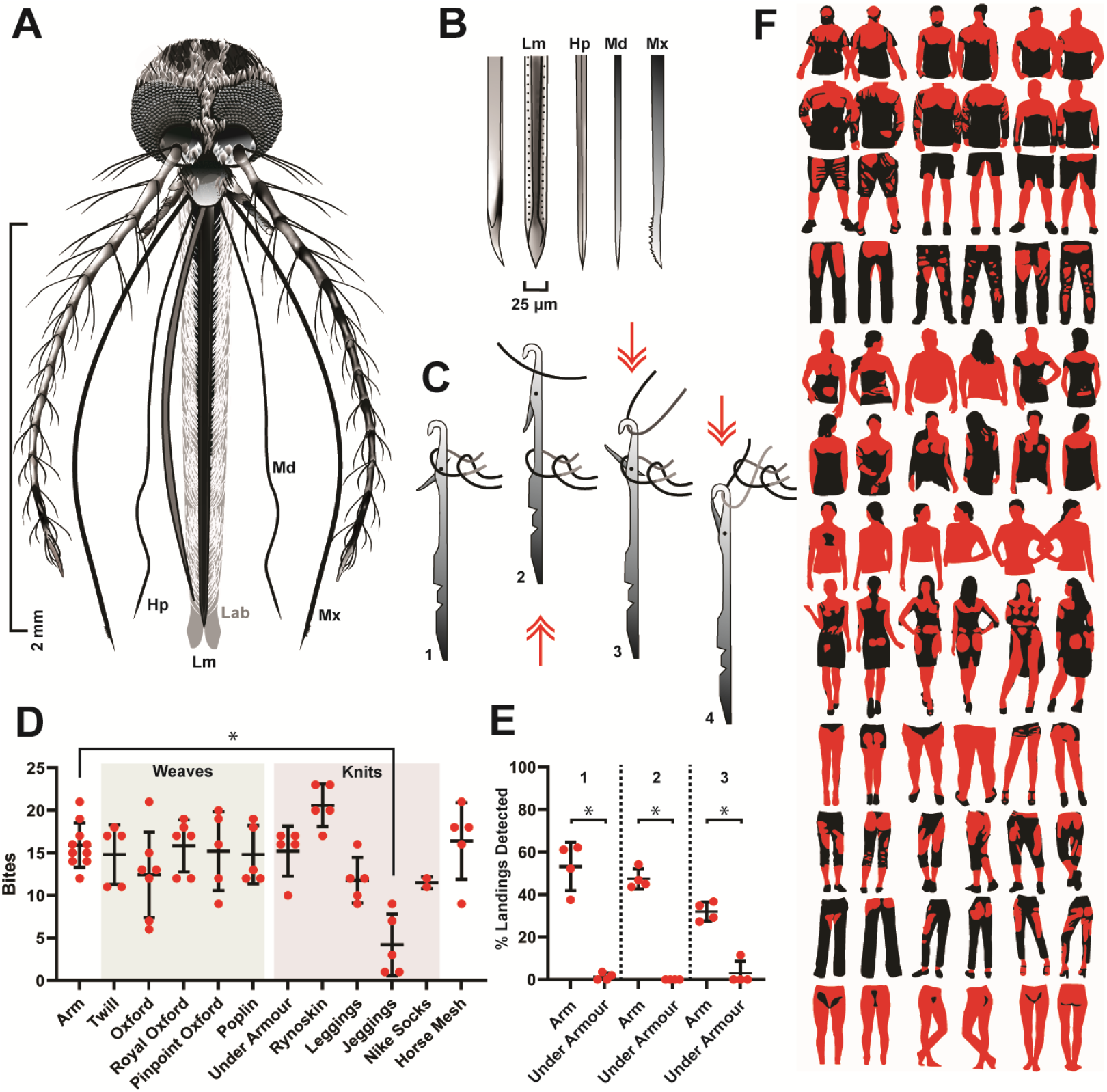
**(A)** Mosquito head and mouthparts. **(B)** Enumeration and dimensions of the mosquito fascicle’s microneedles. **(C)** Needles of flatbed knitting machines with order of movement numbered. **(D)** Screen of common textile’s ability to block mosquitos, comparing weave and knit sleeves. Each dot represents data from one replicate experiment with a cage of 20 unique females. Number of bites are quantified on the y-axis. **(E)** Quantification of detected landing events on a sleeved arm. **(F)** Pixel quantification of garment bite risk, defined as where either clothing clings to skin or skin is uncovered (red). Asterisk (*) indicates significance (p<0.05) by ANOVA in (D) or T-Test in (E). Graphs plot mean and standard deviation.

Modern clothing is manufactured as weaves or knits (**Fig S1**). Weaving interlaces multiple weft and warp fibers, whereas knitting constructs recursive loops from a single fiber forming courses (rows) and wales (columns). Unique knit patterns can be described through symbol diagrams that convey the knit geometry, termed knit diagrams (**Fig S1C-E**) ^25–27^. The written knit code can be translated into robotic machine primitives interpretable by modern flatbed computer numerical control (CNC) knitting machines, which can knit complicated structures with simple up-down needle movements (**Fig 1C**). There are nearly infinite knit configurations and fiber inputs. Thus, the search space of possible knit permutations is vast. Our hypothesis was simply that certain textile configurations would block mosquito bites and others would not. We sought to test and define them.

We performed an initial blocking screen on common clothing. Experiments consisted of placing an arm with sleeve in a cage of 20 female mosquitos for 15 minutes. We quantified the number of bites received. Five common weaves tested did not block, but one knit did (**Fig 1D** and **Fig S2**). Notably modern clothing including *Under Armour* compression heat gear, *Nike* socks, and two garments that advertise insect protection including *Rynoskin* and a protective horse mesh, did not block. Microscopy revealed that these textiles were full of spaces through which mosquitos could probe (**Fig S2**). *Under Armour* also reduced perception of mosquito landing events (**Fig 1E**); as such it’s worse than exposed bare skin. Mosquitos easily pierce clothes, so we quantified where clothing clings to skin for the most common garments in males and females (red in **Fig 1F**). Notably, protection afforded from long sleeves isn’t much better than that of short sleeves because both cling to skin in large areas of the upper back, shoulders, and on parts of the arms (**Fig 1F**). How clothing fit individual bodies is also a factor in getting bit.

Results from initial experiments indicated that knits were capable of blocking mosquito bites with variable efficacy. We sought to determine which features and parameters created the blocking effect. We then screened eight distinct knit geometries (**Fig 2, Fig S3**). We also simulated these knit geometries to facilitate geometric comprehensibility (**Fig S3**). We observed that heat from a standard wash-dry cycle shrunk polyester knits. In the interlock knit, post-knit heat treatment converted a non-blocker into a blocker by shrinking inter-wale and inter-loop space (**Fig I-J**). Thereafter, all knits we tested were heat treated via a wash/dry cycle. Of eight knits screened only one blocked, which was interlock. The interlock knit uniquely positions interlocking loops on top of each other (**Fig 2D**).

**Figure 2.**
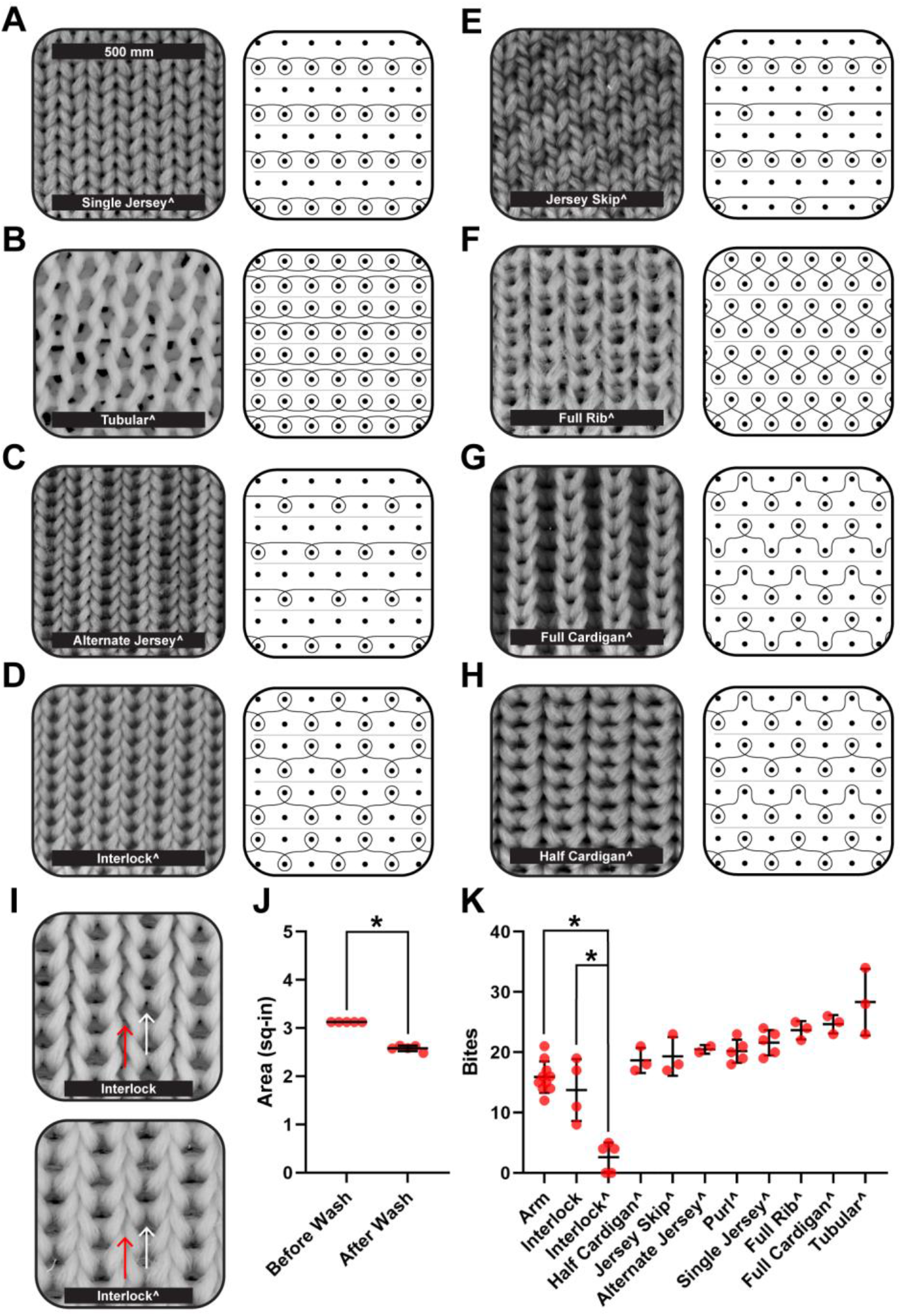
Bite blocking data of all knitted textiles developed at Auburn University using the 100% polyester control yarn of 282 microns. (**A-H**) Microscopy images and knit diagrams of each knit developed. (**I**) Microscopy images of an unwashed (top) and washed (bottom) interlock knit. Microscopy images are zoomed in at a greater scale. Arrows are used to highlight significant points of shrinking which enhance the knits bite blocking abilities. (**J**) Graph of total surface area of an interlock textile measured before and after washing. This experiment was conducted 5 times, each dot representing a single interlock piece that was washed, dried, and then measured. (**K**) Graph of the number of bites received during a single 15-minute experiment. Each red dot corresponds to one cage of twenty females and each sleeve was tested a minimum of three times. Carrot (^) indicates washed. Asterisk (*) indicates p<0.05. Graphs plot mean and standard deviation.

Because we were able to convert a non-blocking knit into a blocking knit, we sought to define treatments and parameters that improve blocking. We discovered three other parameters capable of enhancing mosquito blocking. Increasing thread diameter converted a single-jersey knit into a blocking knit (**Fig 3A**). Increasing spandex content converted jersey-skip knits to blockers (**Fig 3B**). Finally, decreasing stitch length enhanced blocking of the interlock knit (**Fig 3C**). When mosquitos land on blocking knits they tend to probe more, though total probing time was less because they fly away when they cannot get bloodmeals (**Fig 3D**). We tested our blocking knits against both *Aedes aegypti* and *Psorophora howardii. Psorophora* is colloquially known as a “giant mosquito” and its proboscis is much larger than *Aedes* (**Fig S4**). None of the textiles tested were thicker than mosquito proboscises (**Fig S5**). This shows that our knits block more than one species of mosquito. We conclude that unique recipes and optimizations are often required to generate the blocking effect. Moreover, programmed knit diagrams often come out geometrically different postproduction because compressive forces alter the structures, as observed with interlock.

**Figure 3.**
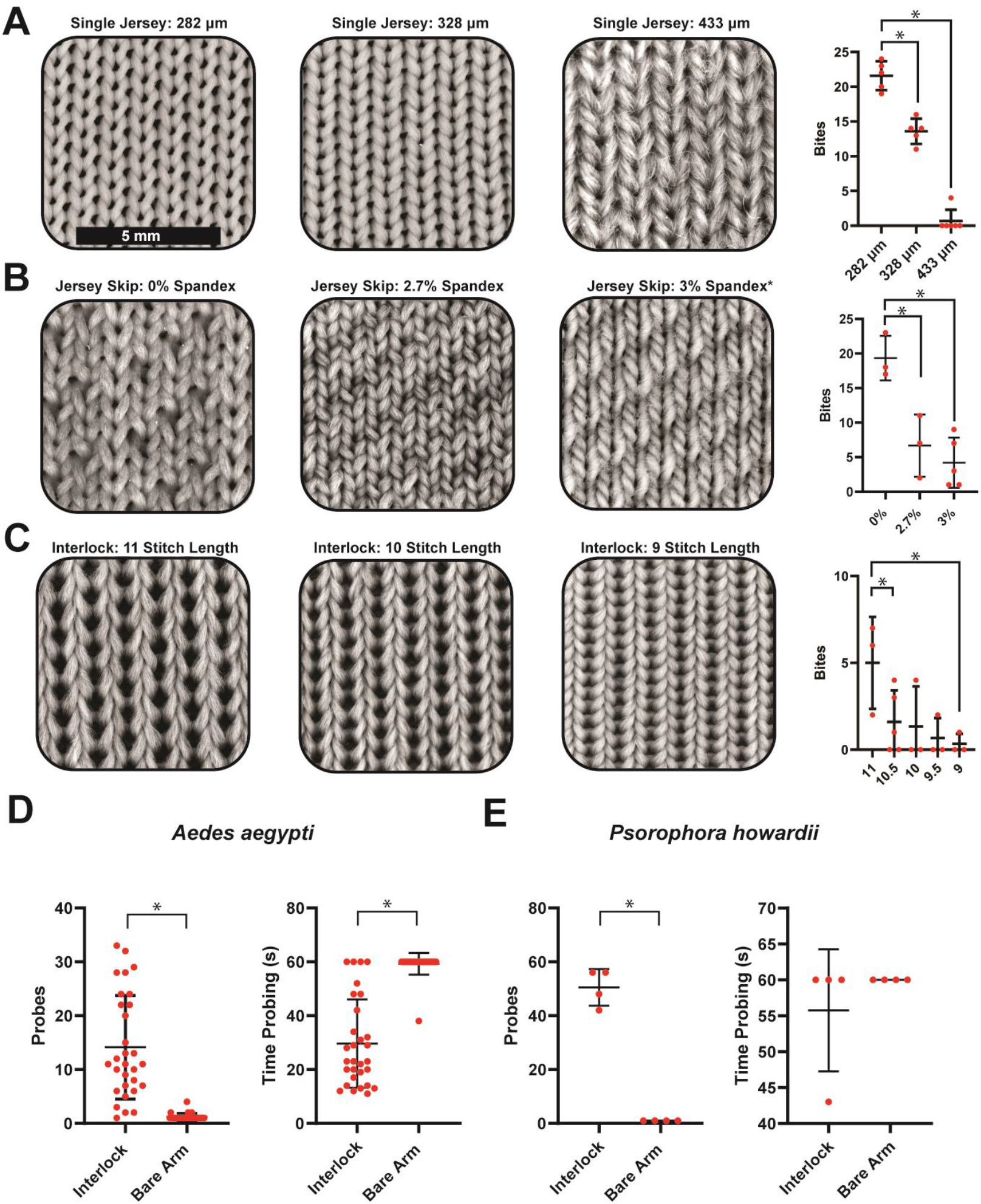
**(A)** Increasing fiber diameter enhances bite blocking in single jersey knits. Numbers above textiles indicate fiber diameter in μm. Scale bar is 5 mm for all images. As before, red dots are independent mosquito bite experiments with 20 females. Graphs show mean and standard deviation. **(B)** Increasing spandex content compresses knit conformations and enhances bite blocking in jersey-skip knits. The final panel (3% spandex*) is constructed of 3% spandex, 19% polyester, and 78% cotton. **(C)** Decreasing stitch length enhanced bite blocking in interlock knits. (D) Quantification of *Aedes aegypti* probing on interlock (stitch length - 10) vs bare arm. Red dots indicate number of probes from an individual mosquito (left); and time spent probing in seconds (s) for individual mosquitos (right). Graphs plot mean and standard deviation. **(E)** Similar probing experiments with *Psorophera howardii* on interlock (stitch length - 10) vs bare arm. Graphs are same as D. Asterisk (*) indicates significance where p<0.05.

We sought to measure and engineer comfort of blocking textiles (**Fig 4, Fig S6**). We measured comfort by a combination of experiments including a 9-factor comfort score including grittiness, fuzziness, thickness, tensile stretch, hand friction, fabric to fabric friction, force to compress, stiffness, and noise intensity ^28,29^. Our three blocking textiles ranged in comfort (**Fig 4A-D**). We found that blockers were near or even better than the *Under Armour* control in some comfort measures. Interlock was lower in comfort score because of its stiffness, thickness, and lack of stretch (which in turn makes it a better blocker). Further iterations of interlock can incorporate spandex/elastic or alternate fibers to increase comfort. Jersey-skip with spandex were the best blockers and were extremely close in comfort to *Under Armour* due to its mixture of materials (spandex, polyester, and cotton). Mosquitos have feeding/taste preferences ^30^. Because mosquitos target preferential skin areas for biting (**Fig 4E**) textiles might be engineered to include blocking knits in regions highly attractive to mosquitos and looser more comfortable knits in regions unattractive to mosquitos. Garments could also be patterned with colors that are less attractive to mosquitos, like white (**Fig 4F**).

**Figure 4.**
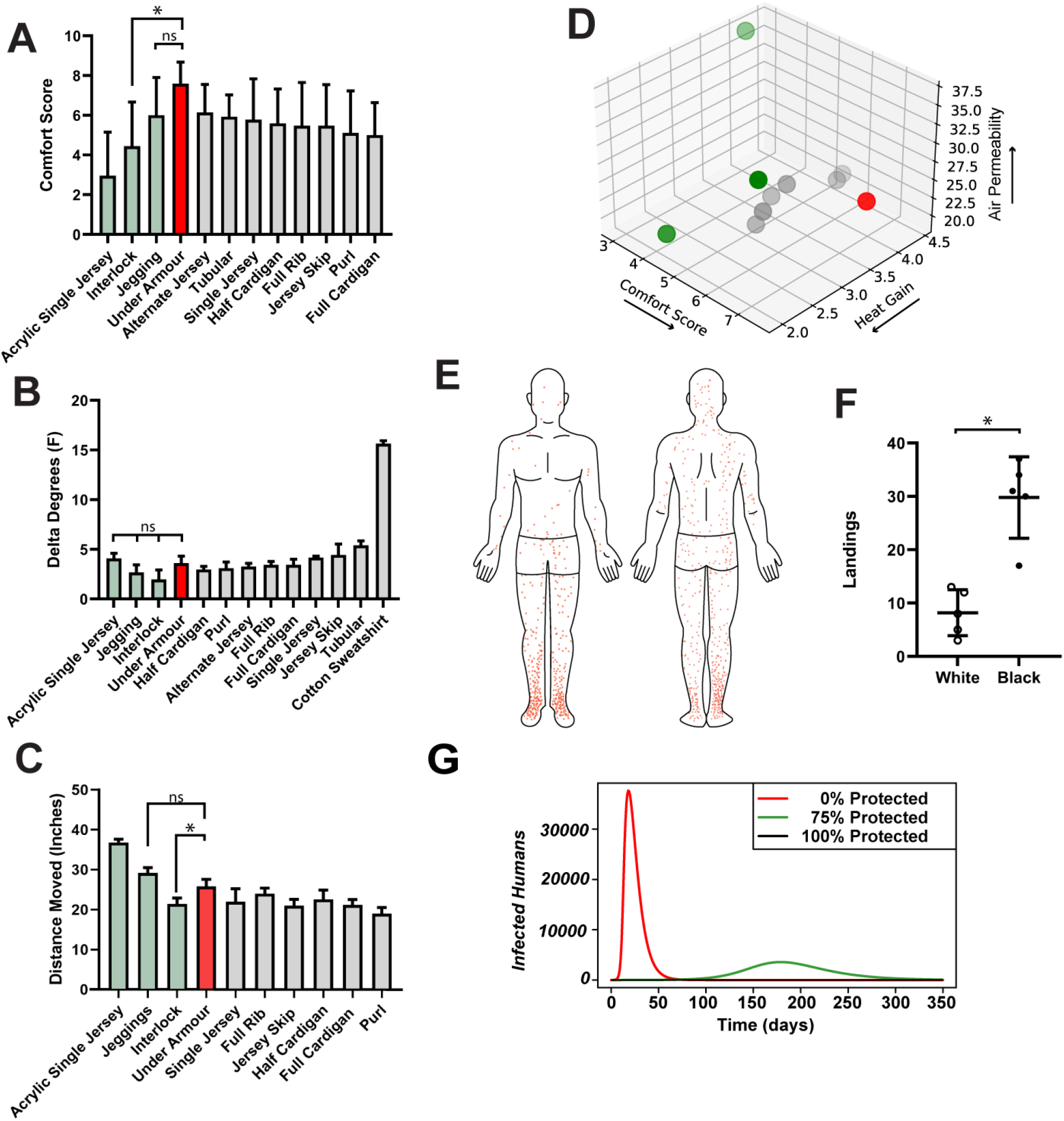
Engineering comfort of bite blocking textiles. **(A)** Mean comfort scores of combined 9-factor feel tests on textiles. A higher score indicates higher comfort. Red is a comfortable *Under Armour* control and green are blocking textiles. **(B)** Average heat gained on skin beneath textile sleeves. A lower score indicates higher comfort (less heat gained). **(C)** Average air permeability of corresponding textile sleeves. Y-axis is distance of salt particles moved by air passing through the textile at 100 psi. A higher score indicates higher comfort (more airflow). **(D)** 3-dimensional comfort graph of textiles. Colors are same as above. Arrow direction indicates increasing comfort.(E) Heat map of mosquito landing events (red dots). Left is front and right is back. **(F)** Choice tests of mosquito landing events on black vs white sleeve regions. Graphs plot mean with standard deviation. Asterisk (*) indicates significance where p<0.05. **(G)** SIR Dengue model over time showing different percentages of individuals protected by bite blocking textiles.

While the immediate benefits of our textiles are clear, we also wanted to understand their potential impact in a real-world outbreak scenario^31^. We simulated a dengue outbreak in a susceptible infected recovered (SIR) model with varying levels of textile protection applied to the population (**Fig 4G**). In the SIR model, as more individuals wear the protective garment, the daily infection rate decreases, delaying and reducing the outbreak peak.

Overall, we showed that modern comfortable textiles can be engineered to block mosquito bites. Blocking sometimes comes at the cost of comfort but needn’t. Our discoveries arm individuals with the power to protect themselves from vector-borne disease in hot climates. The manufacturing process of these textile garments reduces human labor and will not negatively impact the environment.

## Data and Materials Availability

All data are available in the main text or the supplementary materials.

## Materials and Methods

### Knitting Garments

We used M1 Plus (Stoll) to design knit pattern files. Pattern files were loaded into a ADF 530-16 Ki BcW flatbed knitting machines (Stoll, Reutlingen, Baden-Wurttemberg, Germany). Knitted sheets were cut and sewn using as CS7000X sewing machine (Brother, Bridgewater, New Jersey) into sleeves to fit comfortably on the experimenter’s arm. Next hems are created, and raw edges finished by a 1034D serger machine (Brother, Bridgewater, New Jersey). A standard control yarn was used for all knits in **Fig 2**. The control yarn is a 100% polyester of size 2/150/96 (number of plies/denier of each ply/number of filaments in each ply) with diameter 282 microns (Unifi, Greensboro, NC). Other yarns used in the development of textiles throughout the course of this research were also acquired from Unifi. Thickness (mm) of each knitted textile was measured with electronic calipers.

### Mosquitos

*Aedes aegypti* (Linnaeus, Rockefeller strain) mosquitos were reared in-house within a pathogen-free insectary. Mosquitos were kept at 28°C with a rotating 12-hour light/dark cycle. Mosquito eggs are hatched by submerging egg papers in medium sized shoebox tubs until pupation. Larvae and pupae are fed approximately 5 mL of a baker’s yeast and water mixture. Pupae are transferred by hand into a mesh cage for eclosion. As a control for age, pupae are allowed to eclose for 72 hours, then removed and placed into a new cage; this ensures all mosquitos within a cage are within 3 days age of each other. Mosquito biting experiments were performed with females aged 4-7 days old. Every mosquito experiment was conducted under the same controlled parameters. The mosquitos were first anesthetized on ice and sorted using a cold block. The females were then placed in a cage with only water, starving them for the 8-12 hours prior to the experiment. To maintain normal mosquito circadian rhythm, all experiments were conducted in the afternoon. For each experiment, an experimenter wears a knitted sleeve and covered their hand with a latex glove. The covered arm was placed in a cage of 20 female mosquitos for 15 minutes. After experiments, both bites and percent blood-fed females were recorded.

For full body tests, 40 females are sorted in the same manner. The experimenter dressed in long white sleeves and pants and stood in a full body cage for 15 minutes. The landing events were recorded by two observers, one in the front and one in the back. Each time a mosquito landed and attempted to probe; a mark was recorded on a human body outline corresponding to that area. This experiment performed three separate replications on three separate cages of 40 mosquitos (see **Fig 4E**). The replicates were digitalized and overlayed in Adobe Illustrator to produce a comprehensive heat map of landing events. Landing events on arms wearing *Under Armour* were similarly tracked by two observers with a small arm cage of 5 female mosquitos and a duration of 15 minutes (see **Fig 1E**). Experiments involving bites to humans were conducted under approved AU IRB Protocol #21-278 FB.

To acquire video captures of mosquito feeding and biting, thirty females *Aedes aegypti* mosquitos were transferred into a cage with two arm openings. The experimenter wore a test sleeve and latex gloves. A Moment Macro Lens V2 iPhone camera attachment was used to acquire microscopic videos of mosquito probing behavior. Each video is recorded, tracking the behavior of a single female for 1 minute. The videos were analyzed, and two data sets are generated: time to fly away and number of probes. Color choice landing events were tracked using half white/half black sleeve. Twenty female *Aedes aegypti* mosquitos were allowed to land and probe on either side. Five iterations of this experiment were conducted, rotating the sleeve incrementally to control for lighting and behavioral predispositions.

*Psorophora howardii* (Coquillett) mosquitos were field collected as larvae from flood water in Alabama and reared to adult stages. Larvae were fed 5ml of liquified fish food solution and supplemented a carnivorous diet of culex larvae. After eclosion, they were sex sorted and setup in video capture experiments in the same manner as *Aedes*, with the exception that only four mosquitos were used per experiment, due to the difficulty of collecting/rearing *Psorophora*.

### Comfort Testing

The hand feel test is a test of perceived sensory comfort. Within this test, nine factors are measured on a scale from one to ten ^28,29^. In each test, two textiles are used as controls that equate to the minimum and maximum values on the scale. The nine factors being tested are gritty, fuzzy, thickness, tensile stretch, hand friction, fabric to fabric friction, force to compress, stiffness, and noise intensity (the sound a fabric makes as it rubs on skin). Each sleeve was tested for all factors three times with different individuals. The tests were blind. All values from all tests can be averaged to generate an average “comfort score” shown in **Fig 4A**. Sleeves with higher comfort scores are intuitively more comfortable to humans.

Thermal heat absorbed and released by a textile were measured by the following experiment. An experimenter would place a sleeve on their arm and enter a 28°C (70% Relative Humidity) incubator for 15 minutes. Four total digital thermometer readings were taken, two before (on skin and sleeve) and two after incubation (on skin and sleeve). The difference in temperatures on skin and sleeve is determined post experiment and these values graphed as heat gained.

Air permeability quantifies how breathable a textile is. To measure air permeability, we pass compressed air at 20 psi through the textile and measure its force on the other side by quantifying its ability to move a substrate (20 g salt) a given distance. Distance of the furthest particle is graphed in **Fig 4C**. This experiment is conducted on a black table incrementally marked in inches. Textiles that allow for more air to pass through are more comfortable.

### Modeling Textile Exposure to Mosquito Bites

To determine regions of clothing where textile closely contacted skin (**Fig 1F**) we acquired black and white images of average type individuals wearing common garments including shorts, t-shirts, pants, dresses, leggings, long-sleeve shirts, and compression shirts. We faux colored pixels where textiles were touching skin red (leaving other areas black) and counted the colored pixel area in Adobe Photoshop using Adobe selection tool and measurement log. Using front and back images in triplicate, a two-dimensional surface area approximation could be determined for areas of body exposed to mosquito bites for each garment type.

### Microscopy

All microscopy images were taken using a Nikon SMZ1270 stereomicroscope with a Nikon DS-Fi3 camera. The diameter of each yarn was determined by taking two microscopy images of a knitted fabric with and without a mm ruler at the exact same scale. Adobe Photoshop was used to create a digital micrometer scale. The thickness of each yarn was measured at least three times using this scale. Scale bars used in microscopy figures were created by using a millimeter ruler during imaging. A scale bar was made in photoshop and added to each figure.

### Statistics

All data was recorded and analyzed using GraphPad Prism 9 or Microsoft excel. Two different statistical analyses were performed. First, for any experiment that yielded more than two sets of results, an Ordinary One-Way ANOVA and Tukey’s multiple comparisons tests were performed for significance. For experiments with only two data sets, an unpaired, nonparametric, Mann-Whitney t-test was performed for significance. Any comparison yielding a P-value less than 0.05 is considered significant.

### Modeling Dengue Transmission

We reconstructed a well-known Dengue SIR model in R^31^. The model assumes that individuals are infected only once and recover with no chance of reinfection. The bite rate factor has been adjusted to reflect individuals wearing protective clothing, with the percentage of the population determining the change in bite rate. The mosquito’s bite rate is influenced by both the variable ‘b’, representing the average number of daily bites a mosquito delivers, and ‘p_value’, which denotes the fraction of the population still susceptible to mosquito bites despite wearing a protective textile. A ‘p_value’ of 0.2 suggests that only 20% of individuals remain vulnerable, meaning 80% are effectively shielded by the textile. When a mosquito attempts to bite an individual, the combined effect of ‘b’ and ‘p_value’ determines the likelihood of successful transmission. In scenarios where the mosquito targets someone protected, its attempt to spread the disease fails, highlighting the textile’s protective role within the model’s simulated environment.

## Acknowledgments

We thank Stoll and Straehle + Hess for providing knitting expertise. Funding was provided by Alabama Department of Economic and Community Affairs (ADECA) grant 1ARDEF20 02 (JB) and USDA Hatch Grant 1015922 (JB) and Auburn University startup funds (JB).

## Author Contributions

Conceptualization: JB Methodology: All authors Investigation: All authors Visualization: JB, JH, KO Funding acquisition: JB Project administration: JB Supervision: JB, SA, JM Writing – original draft: JB, JH, KO Writing – review & editing: All authors

## Supplementary Materials

Supplementary Information is available for this paper

## List of Supplemental Materials

### Materials and Methods

Fig. S1.

Fig. S2.

Fig. S3.

Fig. S4.

Fig. S5.

Fig. S6.

**Figure S1.**
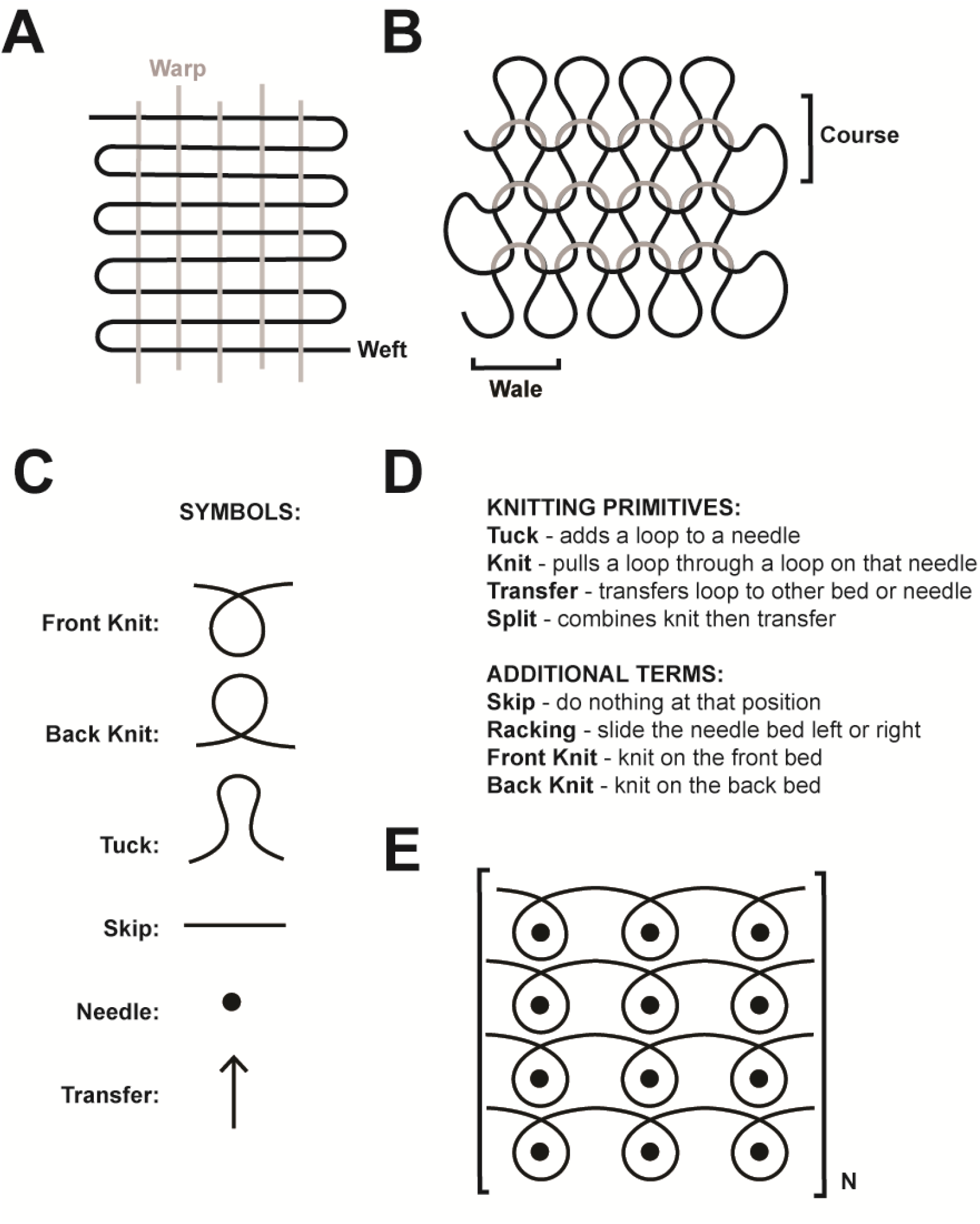
**(A)** Process of weave manufacturing uses multiple independent warp and weft fibers. **(B)** Process of knit manufacturing uses single fibers and draws loops through loops to form courses (rows) and whales (columns) which form a sheet. **(C)** Symbology used in knit diagrams. Front knits are pulling a loop onto the front needle bed and a back knit onto the back needle bed. Tuck draws a loop onto a needle but doesn’t pull it through a prior loop. Skip passes a needle. Dots represent needles and arrows represent transfers of loops from back to front needle bed or visa versa. **(D)** Primitive movements of knitting machines are described. **(E)** Represents an example of a knit diagram demonstrating a repeating pattern of front knits. When knit patterns include both front and back beds, two rows of needles are depicted for each course.

**Supplementary Figure 2.**
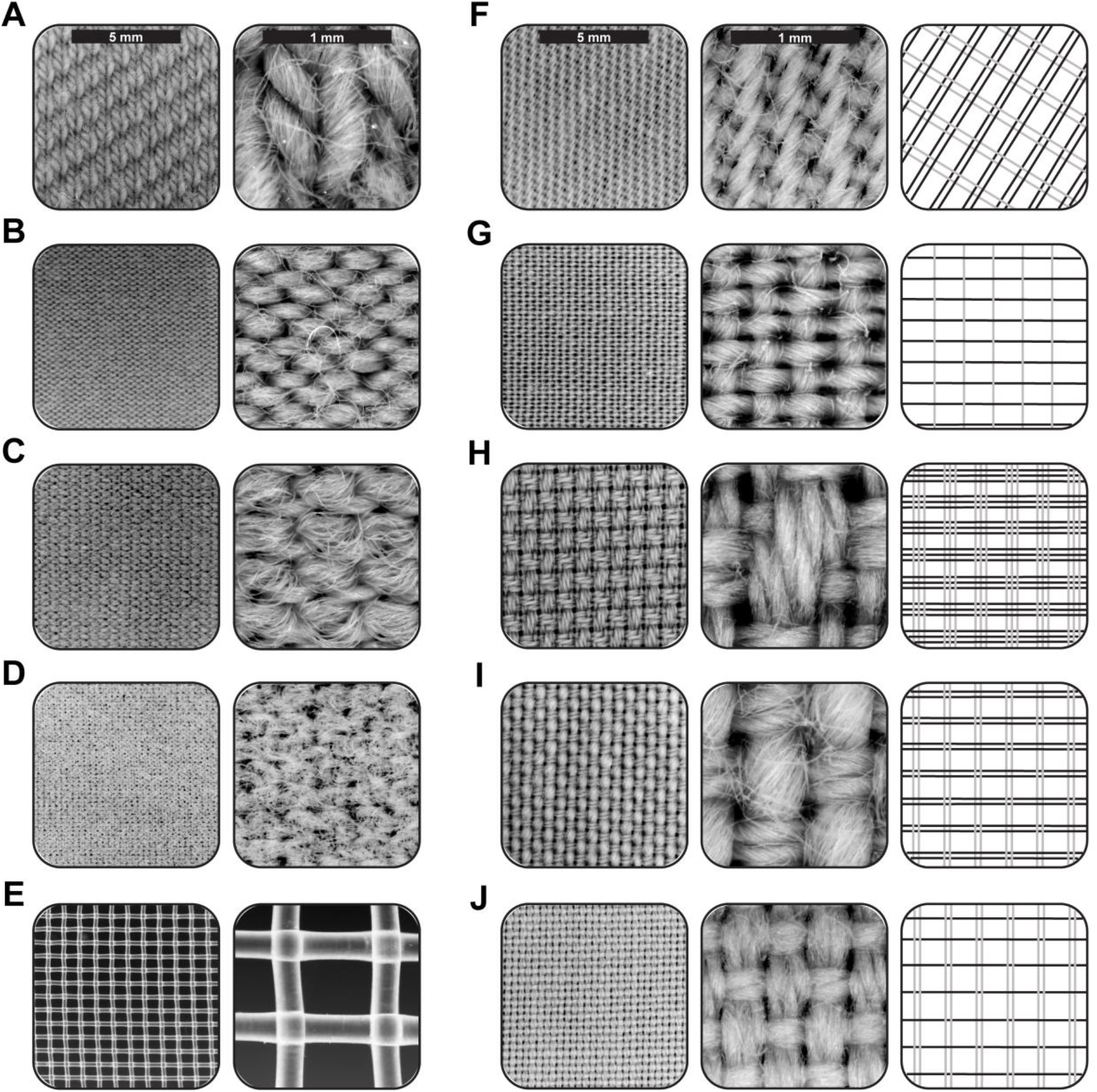
Microscopy of common weaves and knits found in commercial clothing. (**A**) Jegging Weft Knit. (**B**) Legging Weft Knit. (**C**) *Under Armour* Warp Knit. (**D**) Rynoskin Weft Knit. (**E**) Horse Mesh. (**F**) Twill. (**G**) Poplin. (**H**) Royal Oxford. (**I**) Oxford. (**J**) Pinpoint Oxford.

**Supplemental Figure 3.**
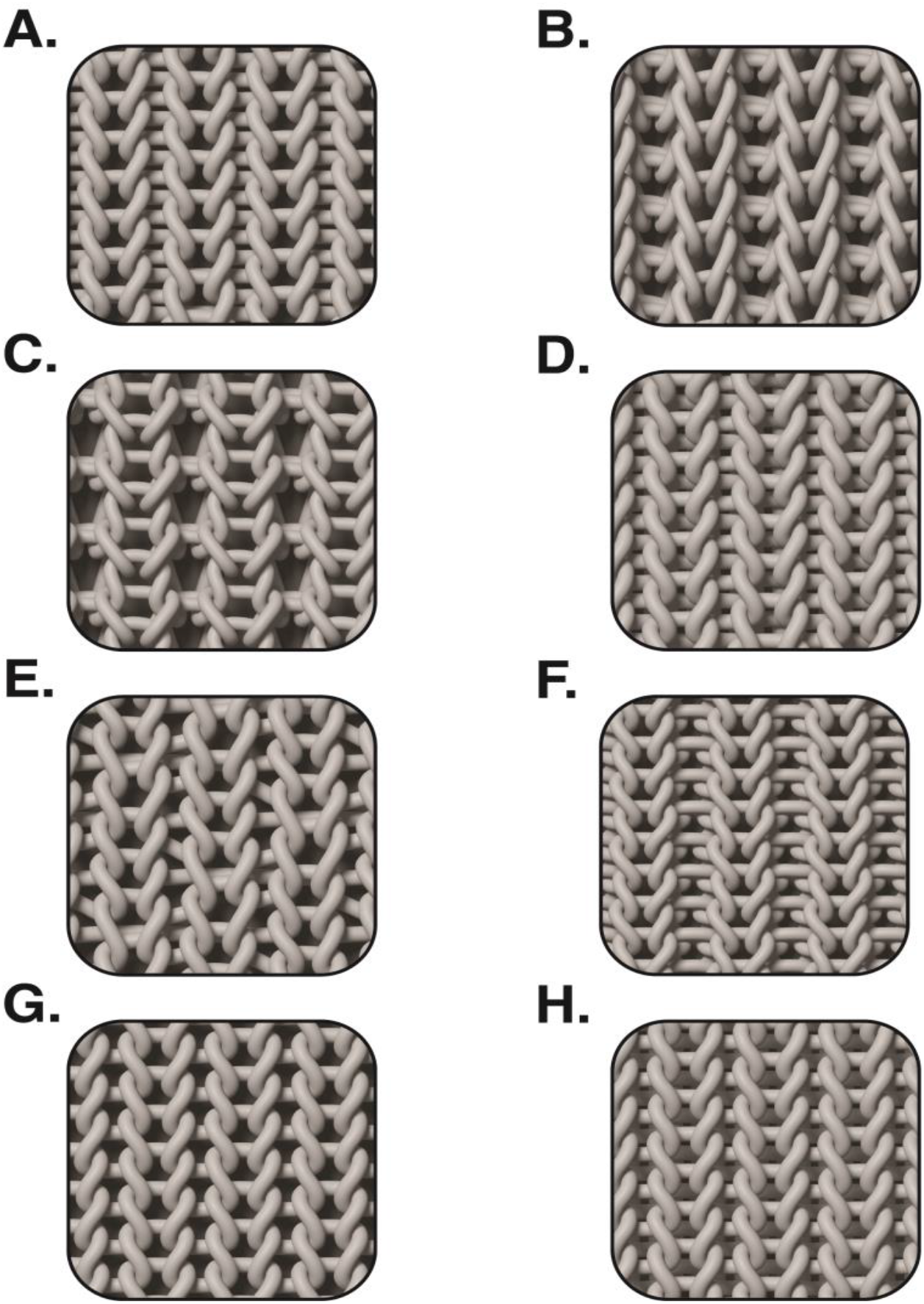
In-silico simulated graphics of knit diagrams facilitate comprehensibility of the knit geometry. **A**. Alternate Jersey. **B**. Full Cardigan. **C**. Half Cardigan. **D**. Interlock. **E**. Jersey-Skip. **F**. Rib **G**. Single Jersey. **H**. Tubular.

**Supplementary Figure 4.**
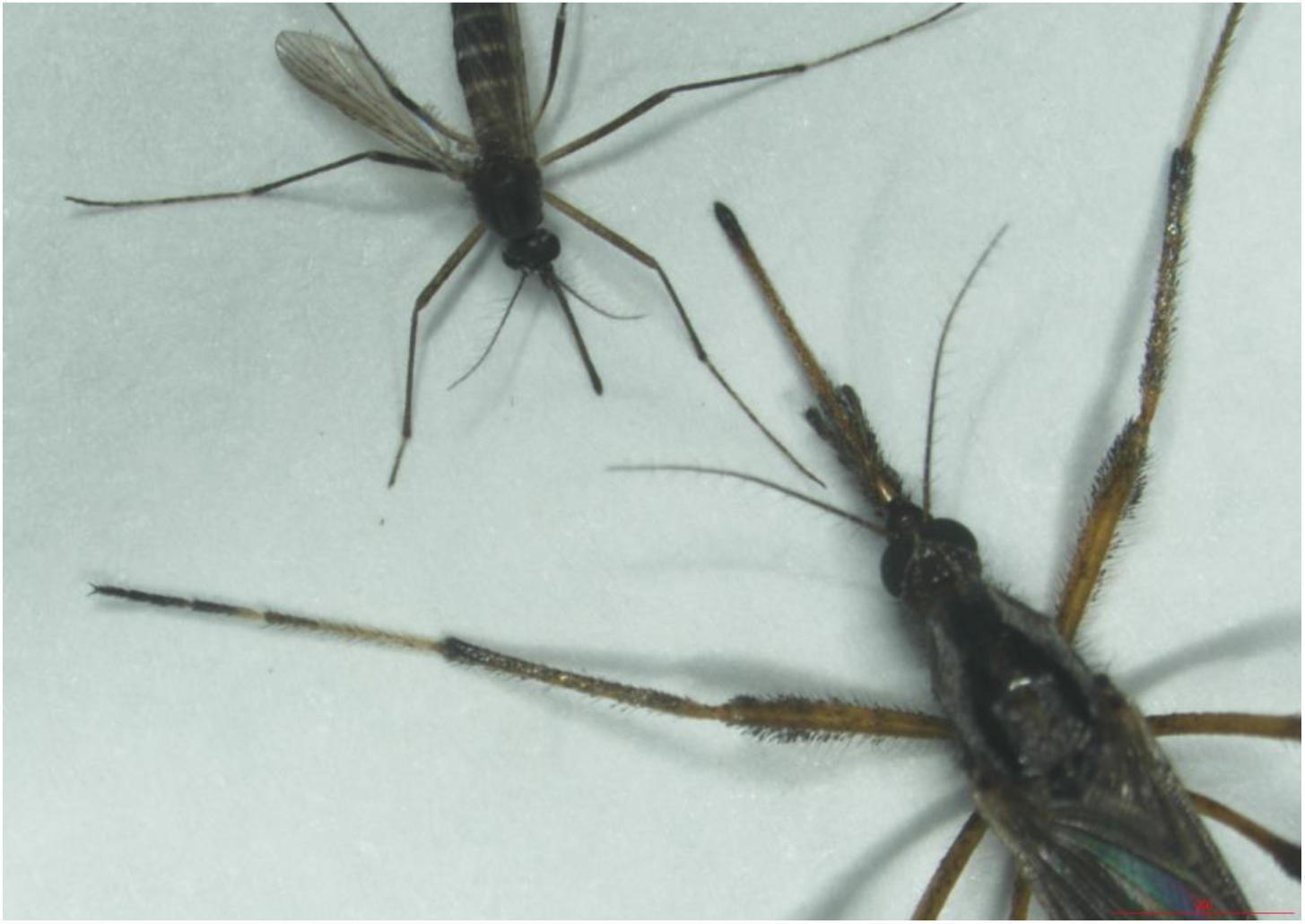
Comparison of *Aedes aegypti* mosquito body and mouthparts against *Psorophora howardii*.

**Supplementary Figure 5.**
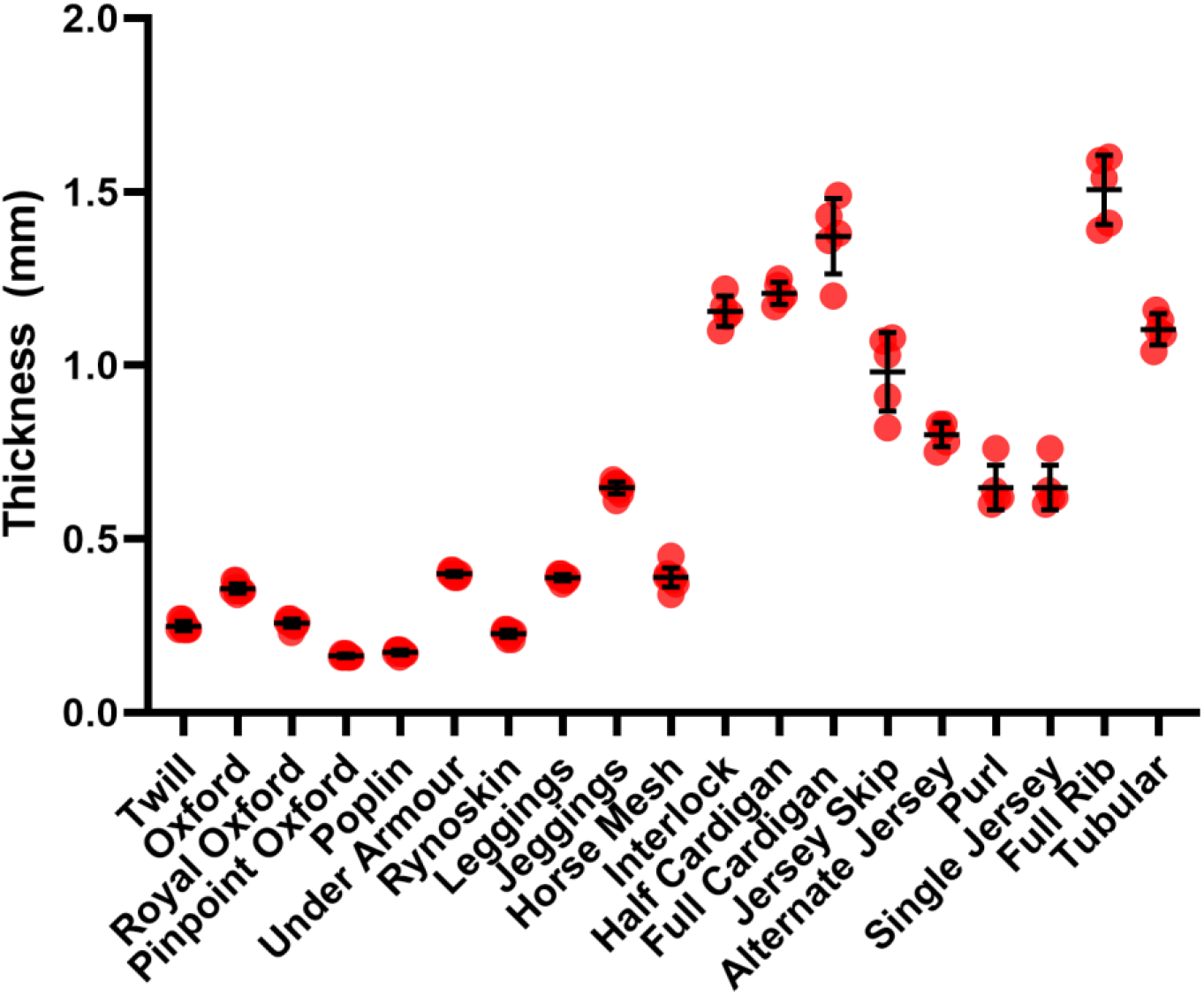
Thicknesses of textiles tested as measured by digital calipers. No textile tested is thicker than the length of an *Aedes* mosquito proboscis, which is 2 mm.

**Supplementary Figure 6.**
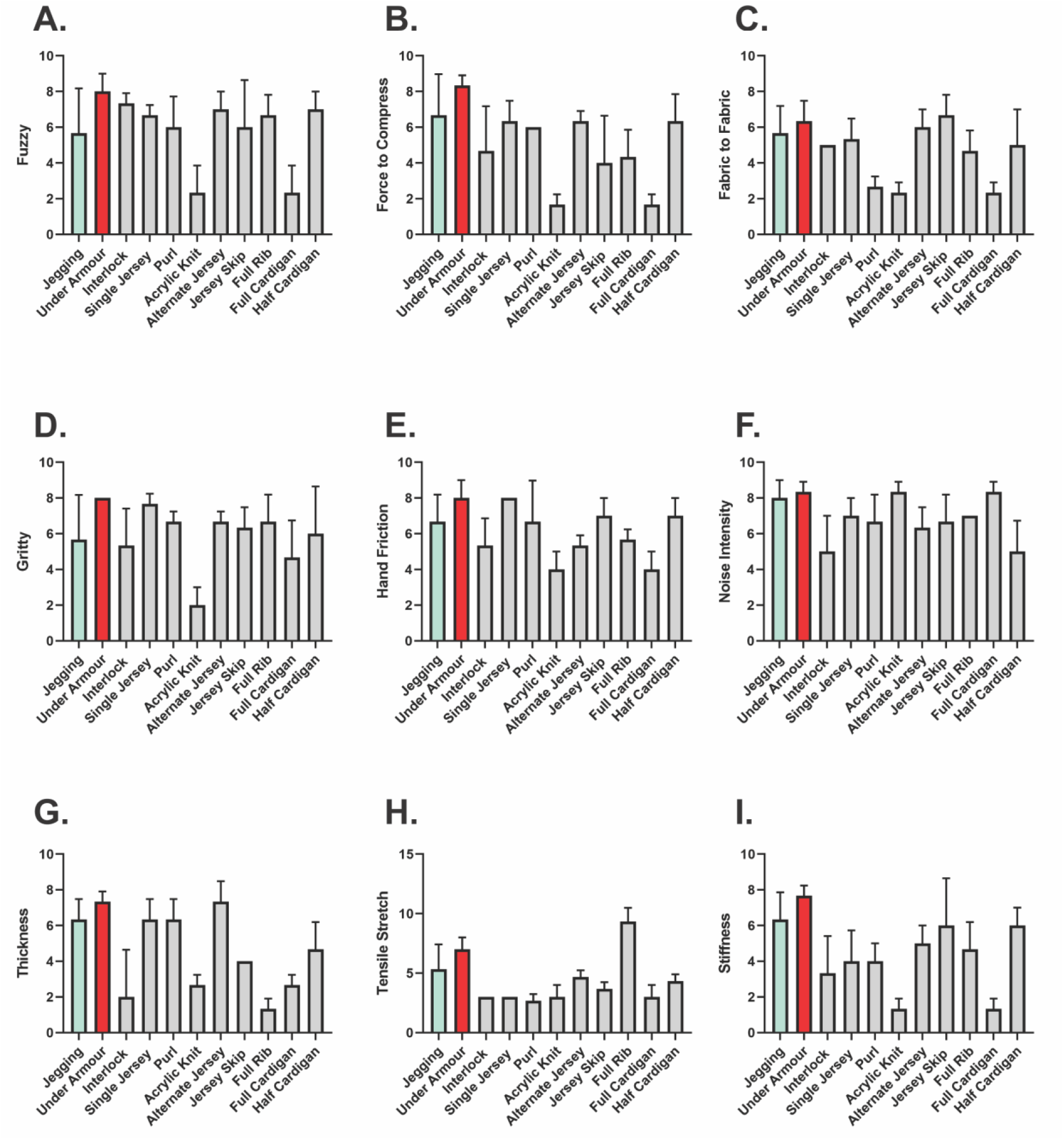
Discrete data from 9 factor determinants of comfort. These data were used to compile a combinatorial comfort score (**Fig 4A**). Graphs are mean with standard deviation. Increased score shows increasing comfort. **A**. Perceived fuzziness **B**. Perceived force required to compress the fabric. **C**. Perceived fabric to fabric friction. **D**. Perceived grittiness. **E**. Perceived hand friction. **F**. Perceived noise intensity (the sound a fabric makes as it rubs on skin). **G**. Perceived thickness. **H**. Perceived tensile stretch. **I**. Perceived stiffness.

